# Structural Basis for Antibody Neutralization of Pertussis Toxin

**DOI:** 10.1101/2024.09.23.614357

**Authors:** Jory A. Goldsmith, Annalee W. Nguyen, Rebecca E. Wilen, Wassana Wijagkanalan, Jason S. McLellan, Jennifer A. Maynard

**Affiliations:** Department of Molecular Biosciences, The University of Texas at Austin, Austin, Texas, USA 78712; Department of Chemical Engineering, The University of Texas at Austin, Austin, Texas, USA 78712; BioNet Asia Co. Ltd., Prakanong, Bangkok, Thailand

## Abstract

Pertussis toxin (PT) is a key protective antigen in vaccine- and natural immunity-mediated protection from *Bordetella pertussis* infection. Despite its importance, no PT-neutralizing epitopes have been characterized structurally. To define neutralizing epitopes and identify key structural elements to preserve during PT antigen design, we determined a 3.6 Å cryo-electron microscopy structure of genetically detoxified PT (PTg) bound to hu11E6 and hu1B7, two potently neutralizing anti-PT antibodies with complementary mechanisms: disruption of toxin adhesion to cells and intracellular activities, respectively. Hu11E6 bound the paralogous S2 and S3 subunits of PTg via a conserved epitope, but surprisingly did not span the sialic acid binding site implicated in toxin adhesion. High-throughput glycan array analysis showed that hu11E6 specifically prevents PTg binding to sialylated N-glycans, while a T cell activation assay showed that hu11E6 blocks PTg mitogenic activities to define the neutralizing mechanism. Hu1B7 bound a quaternary epitope spanning the S1 and S5 subunits, although functional studies of hu1B7 variants suggested that S5 binding is not involved in its PT neutralization mechanism. These results are the first to structurally define neutralizing epitopes on PT, improving our molecular understanding of immune protection from *B. pertussis* and providing key information for the future development of PT immunogens.

**SIGNIFICANCE:** Antibodies neutralizing pertussis toxin (PT) prevent the severe clinical symptoms associated with infection by *Bordetella pertussis*. However, the molecular basis of effective PT-targeted immunity is poorly understood. To gain insight into PT-inhibitory mechanisms, we determined the cryo-electron microscopy structure of genetically detoxified PT (PTg) with two potently neutralizing antibodies to precisely define their epitopes. Carbohydrate-binding studies show that the hu11E6-binding surface on PT interacts with N-linked glycans and that blocking these interactions prevents PT’s T cell mitogenic activities. Hu1B7 binds an epitope near the S1 active site that includes S5 contacts but these do not appear important for neutralization. This work identifies PT-neutralizing epitopes and supports inclusion of the hu1B7 and hu11E6 epitopes in next-generation vaccines and PT-based immunogens.

## INTRODUCTION

*Bordetella pertussis,* the common respiratory pathogen and causative agent of whooping cough, continues to be responsible for ∼1% of global childhood mortality despite the wide availability of effective vaccines [1]. Development of inactivated whole-cell pertussis vaccines (wP) decreased the annual U.S. case load from over 150,000 in the 1930s to <5,000 in the 1970s. However, high lot-to-lot variability and reactogenicity following vaccination with wP led to the subsequent development and adoption of acellular vaccines (aP), which contain purified virulence factors such as pertussis toxin (PT), filamentous hemagglutinin (FHA), type 2 and type 3 fimbriae (FIM2/3) and pertactin (PRN) [2].

The rising incidence of *B. pertussis* infection over the past two decades suggests that aP vaccines may be less efficacious than wP vaccines [3]. Outbreaks in adolescents have been observed in populations vaccinated with aP, suggesting that aP-induced immunity wanes rapidly and contributes to the current large outbreaks across the UK and EU [4, 5]. In addition, the upregulation of PT and the loss of PRN observed in clinical isolates suggest that circulating *B. pertussis* may adapt in response to selective pressure imposed by vaccines [6]. To combat pertussis resurgence, improved acellular pertussis vaccines are needed, but little is known about the protective epitopes on any pertussis vaccine antigen.

PT is secreted at high levels by *B. pertussis* during human infection and is a primary component of all licensed acellular pertussis vaccines. Titers of PT-binding antibodies are predictive of recent infection and high anti-PT titers correlate with less severe disease in humans [7, 8], although there is no accepted serological correlate of protection [9]. In addition, experimental infection in the baboon model indicates that PT-neutralizing antibodies are necessary and sufficient to block disease symptoms without altering bacterial colonization [10][11]. Many currently circulating clinical isolates include the PTP3 promoter change that increases PT expression relative to isolates from previous decades [12], indicating PT will continue to be a key target in future vaccines [2].

PT is a classic AB_5_ toxin, with a catalytically active S1 (A) subunit that sits atop a donut-shaped B_5_ subunit. Five structural proteins combine to form the receptor-binding B subunit: the homologous S2 and S3 subunits, which each form heterodimers with S4, and then associate with S5 [13]. Binding to sialylated glycoproteins on the target cell is mediated by carbohydrate-binding regions in the S2 and S3 domains [14, 15] and is followed by receptor-mediated endocytosis and retrograde transport of PT to the ER [16]. Here, the reducing environment and presence of ATP causes S1 dissociation and transport out of the ER by the ER-associated degradation (ERAD) system [17]. Once in the cytoplasm, S1 ADP-ribosylates the alpha subunit of G_i/o_ proteins to disrupt cellular signaling [18]. Phenotypically, PT exerts various systemic effects, including histamine sensitization, islet activation, and leukocytosis [19].

While many studies have examined overall serological responses to PT, fewer have examined the biochemical basis for toxin neutralization. Of the four characterized epitope groups on S1, just one is potently neutralizing in all assays evaluated [20, 21]. Biochemical data suggest the prototypical protective antibody hu1B7 binds an epitope spanning the S1 and B subunits near the active site to interfere with intracellular toxin trafficking [22, 23]. The most protective B subunit antibody characterized, hu11E6, appears to block receptor binding [23]. Since chemical detoxification of PT skews the immune response to non-neutralizing B subunit epitopes [24, 25] and reduces binding by hu1B7, hu11E6 and other neutralizing antibodies [26, 27], genetically detoxified PTg variants such as the R9K and G129E variants [28] and C180 fragment of S1 [29, 30] are attracting interest for next-generation vaccines. However, no structural data are available for the PT-binding antibodies to define epitopes that merit preservation during antigen engineering.

To gain insight into their inhibitory mechanisms, we sought to define the structural basis for PT-binding by neutralizing antibodies. Using cryo-EM, we determined a high-resolution structure of PTg in complex with the hu1B7 and hu11E6 Fabs, which exhibit synergistic protection in multiple murine and baboon models of disease [10, 31, 32]. Delineation of the key epitopes targeted by hu1B7 and hu11E6 will support use of these antibodies as serological probes, allow for tracking of escape mutations in circulating strains and guide the production of next-generation PT vaccines.

## RESULTS

### Cryo-EM structure of the PTg-hu1B7-hu11E6 complex

To better understand the neutralizing mechanisms of hu1B7 and hu11E6, two of the most potent and best-studied neutralizing antibodies against PT, we aimed to determine a cryo-EM structure of the antigen-binding fragments (Fabs) of these antibodies in complex with PT. As one motivation behind newly developed PTg antigens is preservation of neutralizing epitopes, we complexed the hu1B7 and hu11E6 Fabs to a PTg antigen (R9K / E129G), which was produced in an engineered *B. pertussis* strain [33]. The two amino acid changes in PTg do not appear to impact binding of the parental 1B7 and 11E6 antibodies to PT (Fig. S1).

To confirm the homogeneity of the PTg preparation and complex formation, the PTg-hu1B7- hu11E6 complex was analyzed by negative-stain electron microscopy (nsEM). Micrographs of PTg- hu1B7-hu11E6 adsorbed to the carbon layer of CF-400Cu grids and stained with methylamine tungstate exhibited homogeneous particle distribution (Fig. S2A). 2D class-averages of extracted particles confirmed that PTg was binding to 2 or 3 Fab molecules (Fig. S2B).

To prepare samples for cryo-EM analysis, the PTg-hu1B7-hu11E6 complex was applied to UltrAuFoil grids[34], which contain a holey gold film on a gold support, and subsequently blotted and plunge-frozen using a Vitrobot Mark IV. Initial screening of grids prepared in this manner resulted in heterogeneous particle distributions largely composed of aggregates or small particles that were either excess Fab or dissociated complex (Fig. S3A). The particle homogeneity and lack of aggregation observed in the nsEM images suggested that the heterogeneity observed in the cryo-EM images is specific to the blotting and plunge-freezing process, and that the sample may have been denatured by the air-water interface. The inclusion of detergents in cryo-EM samples can mitigate aggregation and other effects of the air-water interface. Accordingly, we froze grids of PTg-hu1B7-hu11E6 containing 0.1% Amphipol but observed a similar heterogeneous mixture of aggregate and dissociated particles on these grids (Fig. S3B), similar to conditions lacking detergent (Fig. S3A). This prompted us to screen 3 additional detergent conditions: 0.02% fluoro-octyl maltoside, 0.4% CHAPS, and 0.1% β-octyl glucoside. Grids prepared with 0.4% CHAPS contained particles too small to be the PTg-hu1B7-hu11E6 complex and were likely to be excess Fab or dissociated components of the complex (Fig. S3C). The grids prepared with 0.04% fluoro-octyl maltoside contained very few particles within the holes, although they appear intact (Fig. S3D). Notably, grids prepared with 0.1% β-octyl glucoside exhibited a homogeneous population of particles, with little to no aggregate present (Fig S3E). This grid was therefore chosen for cryo-EM data collection.

Before collecting a full dataset, *ab initio* reconstruction and homogeneous refinement were performed on particles extracted from a test dataset containing 20 movies. The orientation distribution plot for this reconstruction displayed extreme preferred orientation bias, with almost every particle adopting the same pose (Fig. S3F). To mitigate the preferred orientation bias, the full dataset of 1,500 movies was collected with a stage tilt of 30°. After motion correction, CTF estimation, and particle picking, 1,694,697 particle picks were extracted and subjected to multiple rounds of *ab initio* reconstruction and heterogeneous refinement. A final stack of 316,937 particles was subjected to homogeneous refinement to yield a 3.6 Å resolution reconstruction (Figs. 1A, S4, S5). Using the crystal structure of PT (PDB ID: 1PRT) as a starting model, a model of the PTg-hu1B7-hu11E6 complex was built into the map (Figs. 1B, S6).

**Figure 1.**
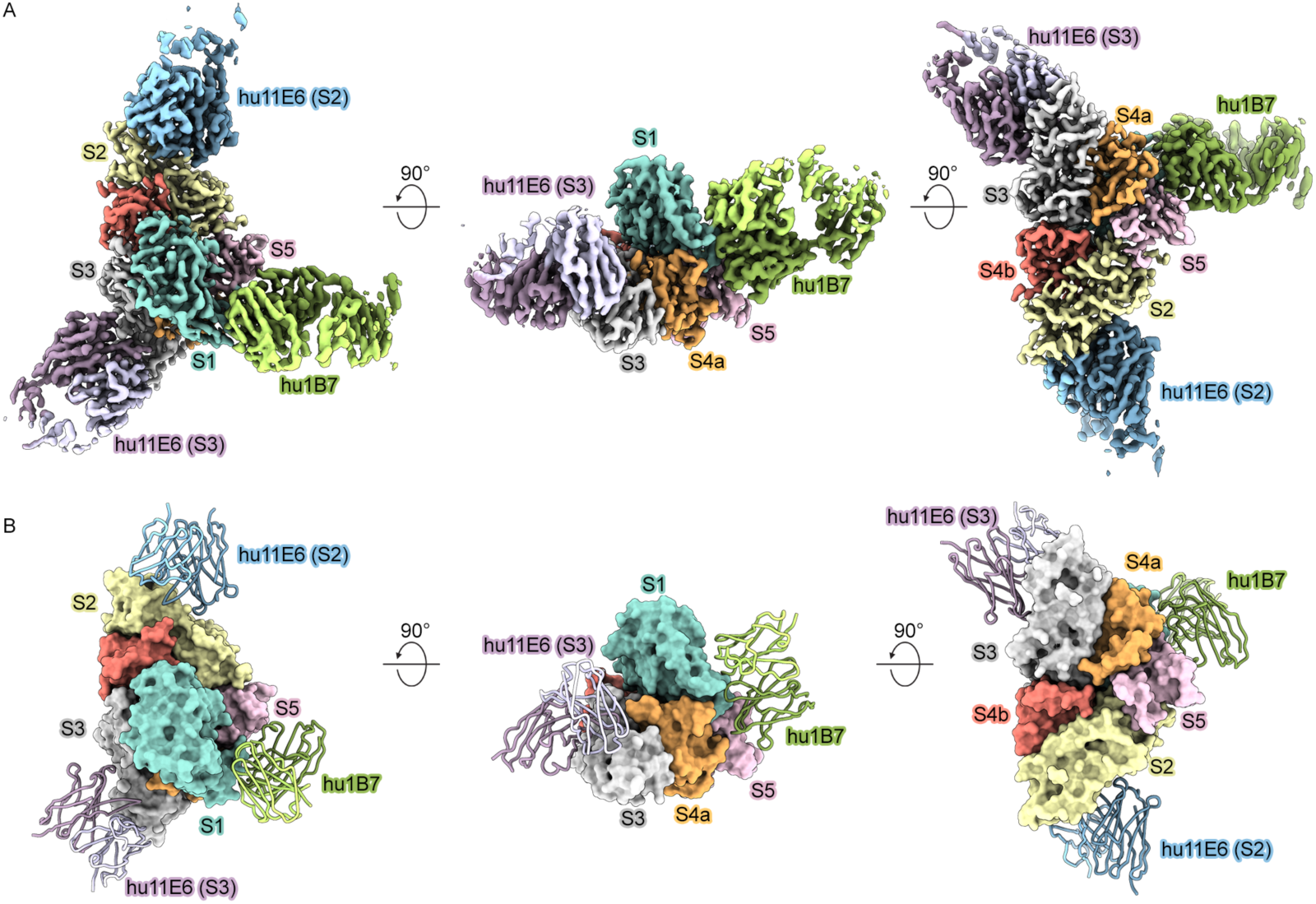
Cryo-electron microscopy structure of PTg in complex with hu1B7 and hu11E6. (A) Sharpened cryo-electron microscopy map of PTg+hu11E6+hu1B7 with S1 colored turquoise, S2 colored yellow, S3 colored white, S4a colored orange, S4b colored red, S5 colored pink, hu11E6 (S2) colored blue, hu11E6 (S3) colored purple, and hu1B7 colored green. (B) Model of PTg+hu11E6+hu1B7 colored as in (A), with PTg subunits shown as molecular surfaces and hu11E6 (S2), hu11E6 (S3) and hu1B7 shown as ribbon representations.

### Hu11E6 binds to a conserved epitope in both S2 and S3 subunits of PT

The paralogous PT S2 and S3 subunits share ∼70% identity across two distinct domains: an N-terminal aerolysin/pertussis toxin domain (APT) with similarities to C-type lectins and a C-terminal B-subunit domain that supports toxin assembly. Despite this modest sequence identity, hu11E6 exhibits a nearly identical binding mode towards S2 and S3 due to hu11E6 binding conserved residues on the APT surface. The light chain (LC) of hu11E6 buries 391 Å^2^ surface area on S3 and 370 Å^2^ surface area on S2, while the heavy chain (HC) of 11E6 buries 453 Å^2^ surface area on S3 and 376 Å^2^ surface area on S2. Ten hydrogen bonds are formed between hu11E6 and S3 (Fig. 2, top), nine of which form in an identical manner between hu11E6 and S2 (Fig. 2, bottom). The only hydrogen bond formed between hu11E6 and S3 that is not formed between hu11E6 and S2 is between the mainchain carbonyl oxygen of hu11E6 HC Asn56 and the sidechain amide nitrogen of Gln182 (Fig. 2, top). In S2, the homologous residue of S3 Gln182 is Glu182, which cannot form the same hydrogen bond with the mainchain carbonyl oxygen of hu11E6 HC Asn56.

**Figure 2.**
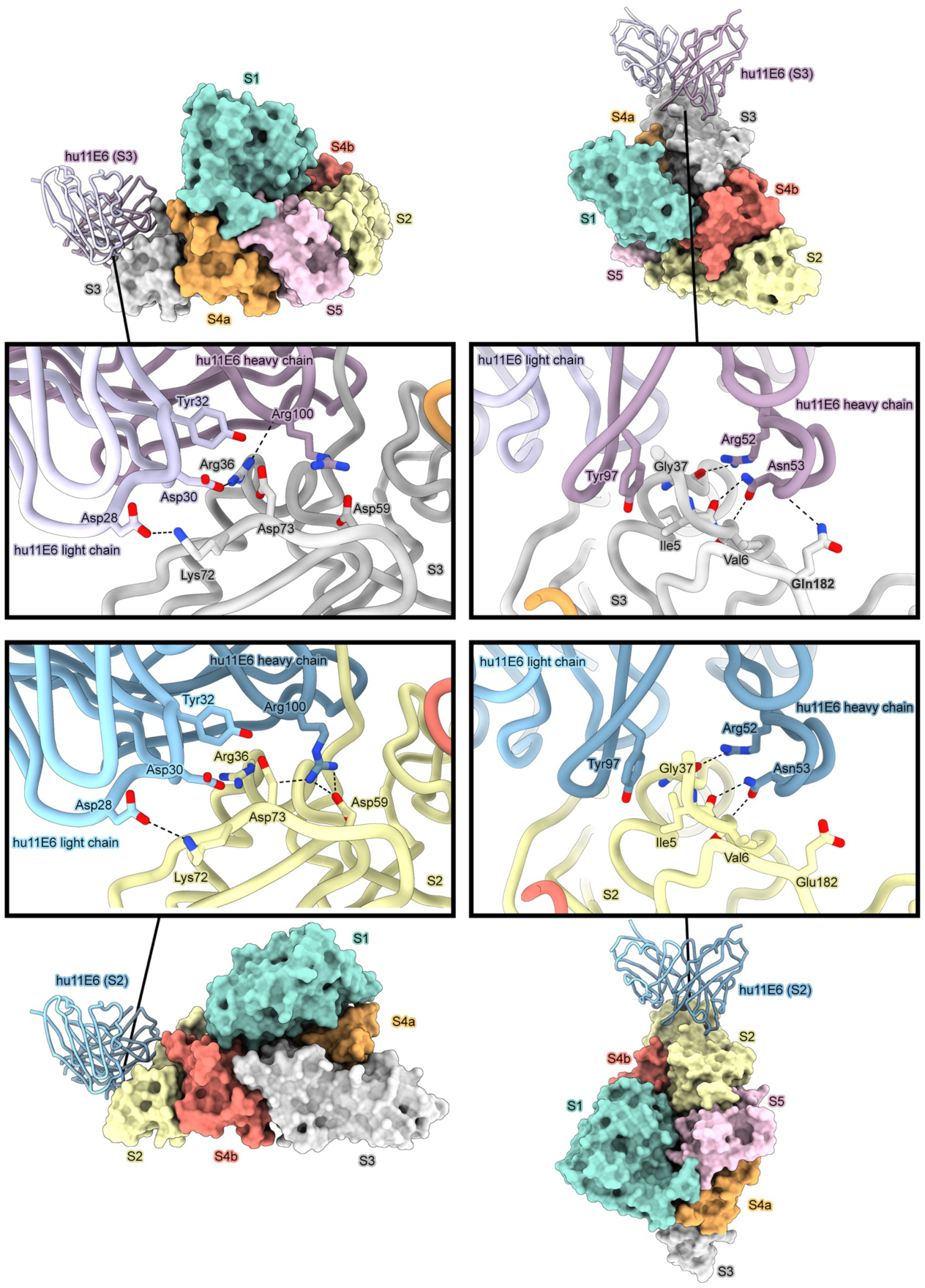
Hu11E6 binds to a conserved epitope on PTg S2 and S3. Zoomed-out views of PTg bound to hu11E6 are shown with PTg as molecular surfaces, with S1 colored turquoise, S2 colored yellow, S3 colored white, S4a colored orange, S4b colored red, S5 colored pink, hu11E6 (S2) colored blue, hu11E6 (S3) colored purple, and hu1B7 colored green. hu11E6 is shown as a ribbon representation. The boxes show close-up views of the hu11E6 epitope on S3 (top) and S2 (bottom) subunits of PTg, with all proteins shown as ribbon representations and key interface residues shown as sticks.

Previously, in an attempt to determine the binding site for its carbohydrate receptor, PT was crystallized in the presence of an 11-mer oligosaccharide, of which the terminal two residues (mannose and sialic acid) were resolved bound to S2 and S3 [35]. Critically, these previously observed sialic acid binding sites do not overlap with the hu11E6-binding sites, but are ∼15–20Å away on the same face of the S2 or S3 subunit (Fig. 3A). It is notable that this face is largely conserved between S2 and S3, with sialic acid and hu11E6 both binding to conserved patches (Fig. 3B). By contrast, the bottom face of S2/S3 (180° opposite the S1 subunit) is formed predominantly by residues that differ between S2 and S3. This suggests that additional carbohydrate-binding sites, which are blocked by hu11E6 binding, are present on the conserved face.

**Figure 3.**
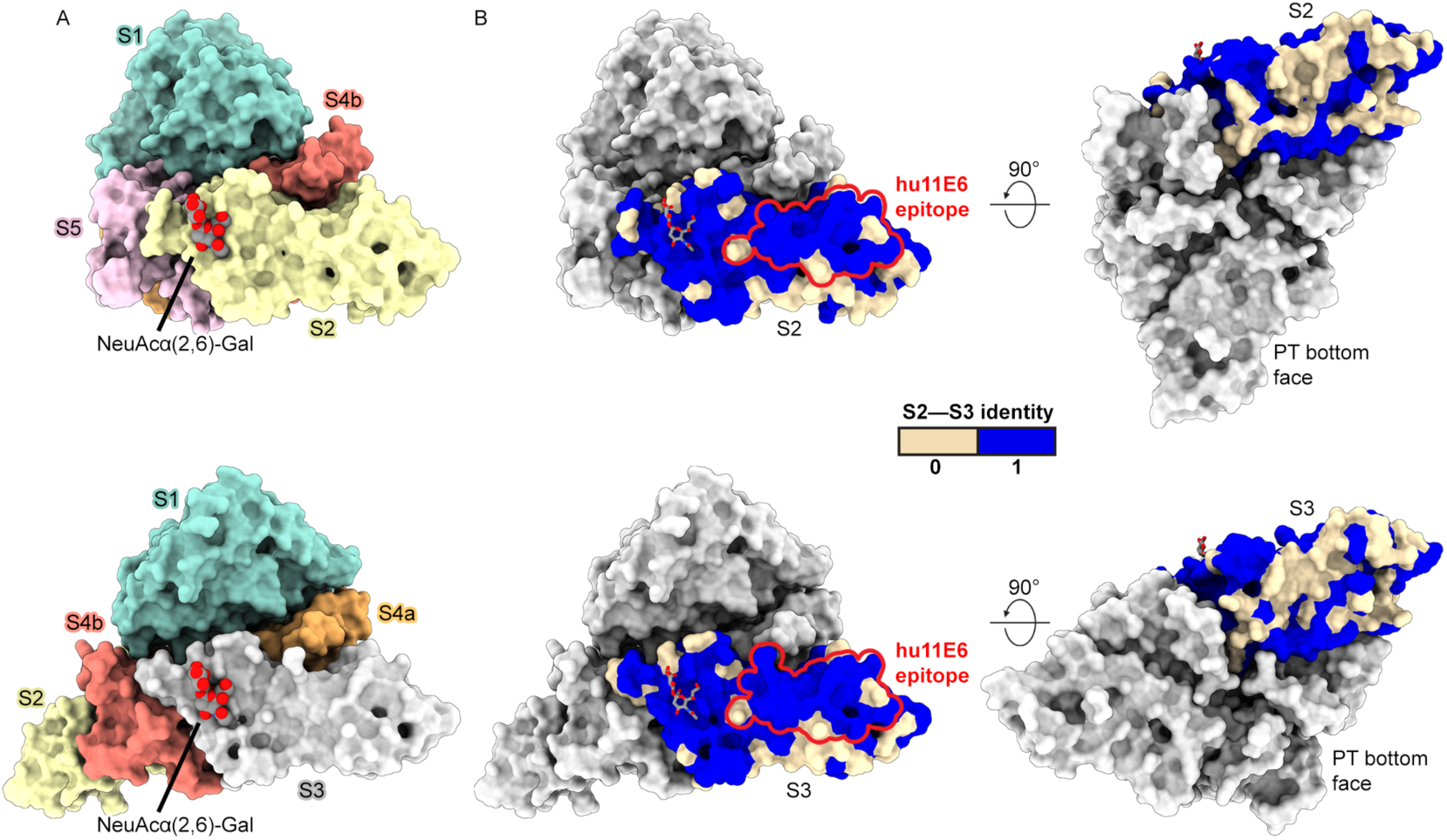
Comparison of hu11E6 epitope and the known sialic acid binding site on PTg. (A) PTg shown as molecular surface with S1 colored turquoise, S2 colored yellow, S3 colored white, S4a colored orange, S4b colored red and S5 colored pink, with NeuAcα(2,6)-Gal disaccharide from PDB ID: 1PTO shown as molecular spheres with carbons colored gray and oxygens colored red. (B) PTg shown as a molecular surface with all subunits colored white and either S2 (left) or S3 (right) colored by per-residue S2–S3 identity, where shared residues between S2 and S3 are colored blue and residues that differ between S2 and S3 are colored tan. The outline of the hu11E6 epitope on S2 and S3 is shown in red. The NeuAcα(2,6)-Gal disaccharide described in (A) is shown on S2 and S3 using stick representation.

### Inhibition of carbohydrate binding by hu11E6

To assess whether hu11E6 directly interferes with glycan recognition, we measured binding of PTg to a high-throughput glycan array containing 562 carbohydrates in the presence and absence of the hu11E6 Fab. PTg alone binds to sialylated N-glycans, asialo-N-glycans and other carbohydrates (such as the sialyl Lewis^X^ blood group antigen (Neu5Acα2-3Galβ1-4(Fucα1-3)GlcNAc) and c-series ganglioside GQ2 (Neu5Acα2-8Neu5Acα2-8Neu5Acα2-8Neu5Acα2-3(GalNAcβ1-4)Galβ1-4Glcβ; Figure 4A-D).

**Figure 4.**
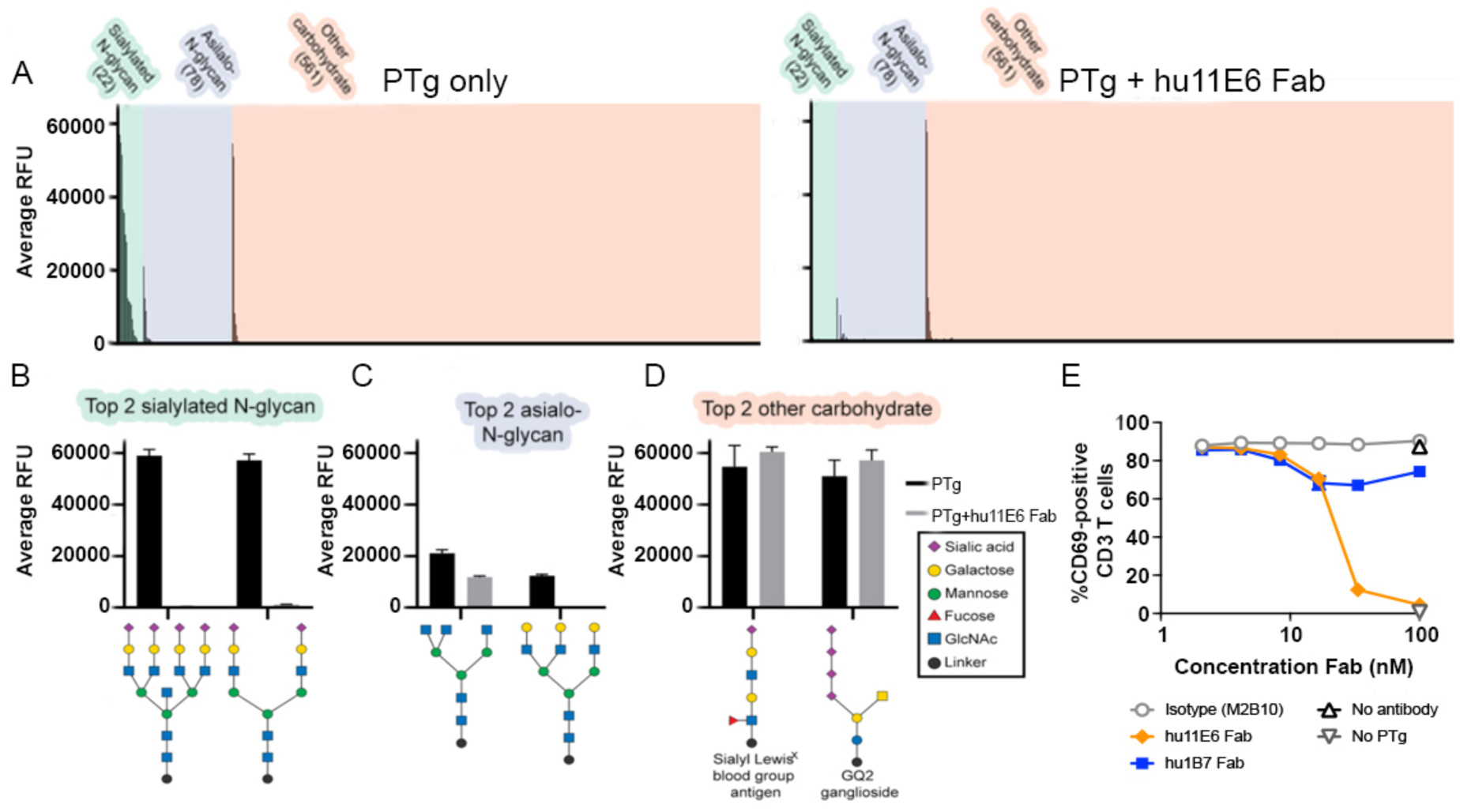
Hu11E6 prevents PTg binding to sialylated N-glycans and T-cell activation. (A) Summary of high-throughput carbohydrate-binding array results for PTg and PTg+hu11E6 Fab. Carbohydrates were separated into 3 groups: sialylated N-glycans (green), asialo-N-glycans (blue), and all other carbohydrates (coral). For PTg (top), carbohydrates were ranked with descending relative fluorescence units (RFU). For PTg+hu11E6 Fab (bottom), carbohydrates were ordered as for PTg alone. Close-up view of the RFU for PTg versus PTg+hu11E6 Fab for the top two (B) sialylated N-glycan hits, (C) asialo-N-glycan hits, and (D) other carbohydrate hits. (E) Antibody Fab inhibition of PTg-mediated T cell activation. Serially diluted Fabs were incubated with PTg and then human donor PBMCs overnight before using flow cytometry to measure the percent of activated, CD69-positive CD3+ T cells. Two-way ANOVA analysis used to determine significance; hu11E6 versus hu1B7 and hu11E6/ hu1B7 versus isotype comparisons indicated with **** reached p<0.0001. Data shown are representative of two biological replicates, with the average and range of two technical replicates shown.

This is similar to a previous glycan array study of PT that demonstrated interactions with 4 major carbohydrates or carbohydrate classes [36]: 1) sialylated N-glycans, 2) asialo-N-glycans, 3) compounds with a terminal Sialo-2,3-lactose, with the strongest hit in this group being sialyl Lewis^X^ and 4) polysialic acid compounds, with the strongest hit in this group being the GQ2. However, when first complexed to hu11E6 Fab, no PTg binding to sialylated N-linked glycans was observed (Figure 4A, 4B). Hu11E6 also reduced binding to some asialo-N-glycans (Figure 4A, 4C) while other carbohydrate hits were unaffected, including the sialyl Lewis^X^ and GQ2 (Figure 4A, 4D). Together, this suggests that hu11E6 blocks an N-linked glycan recognition site in the N-terminal APT domain of S2/S3, whereas additional sites, likely in the C-terminal B-subunit domain, were not blocked.

Previous studies demonstrated PT activation of T cell signaling in an antigen-independent manner leading to T cell anergy, exhaustion, and apoptosis, which is mediated by B-subunit interactions with a receptor thought to be closely associated with or in the TCR complex [37]. To determine whether hu11E6 blockade of PTg–sugar interactions would also diminish T cell activation, we performed an activation assay with PTg, the Fabs of hu1B7, hu11E6 or an isotype control and peripheral blood mononuclear cells (PBMCs) (Fig. 4E). Briefly, PTg was preincubated with serially diluted Fab, then co-incubated overnight with isolated human donor PBMCs, before using flow cytometry to quantify the percent of activated CD3-positive T cells using CD69-upregulation as an activation marker (Fig. S7A). Serially diluted PTg indicated that 9.5 nM induced strong activation and this concentration was selected for antibody inhibition experiments (Fig S7B). Whereas cells incubated with PT and the isotype control showed a high level of T cell activation (∼90% at all antibody concentrations), this was slightly reduced by hu1B7 (∼70% at >10 nM), while hu11E6 showed a nearly complete knockdown of PTg stimulation at higher antibody concentrations (∼5% at 100 nM) and a 50% inhibitory concentration (IC_50_) of 24.5 ±2.5 nM. This indicates that hu11E6 is not only capable of blocking activities mediated by the PT A subunit, but also those mediated by the B subunit [31].

### Hu1B7 binds to a quaternary epitope that spans S1 and S5

In the PTg-hu1B7-hu11E6 complex cryo-EM structure, the hu1B7 heavy chain and light chain bury 428 Å^2^ and 359 Å^2^, respectively, of solvent-accessible surface area on PTg (Fig. 5A, B). Consistent with epitope mapping studies, the hu1B7 light chain (LC) Phe32, Tyr49, Leu50, Trp91 and hu1B7 heavy chain (HC) Ala99 form a hydrophobic interface on the surface of the PTg S1 β-sheet formed by β4, β8 and β9, contacting PT S1 Thr81, His83, Ile85, Tyr148 and Ile152 (Fig. 5C). Similarly, hu1B7 HC Trp33 packs onto the PTg S1 α3–β4 loop, and hu1B7 HC Asn50 and Ser97 form hydrogen bonds with PTg S1 Glu75 and Gly78 within this loop, respectively (Fig. 5D). In addition, hu1B7 heavy chain Ser56 forms a hydrogen-bond between its sidechain oxygen and the C-terminal carboxyl group of S1 Glu99 (Fig. 5D). Biochemical data have previously shown that hu1B7 interacts with PT S1 [24], with residue-level epitope mapping suggesting the involvement of the loops approximately at S1 residues 77–84 (α3–β4 loop) and 149–155 (β8–β9 loop) [22, 38]. C One possible mechanism of inhibition for hu1B7 is the inhibition of S1 catalytic activity. Hu1B7 binding to PTg does not alter the S1 conformation, and the α-carbons of hu1B7-bound S1 align to unbound S1 [13] with an RMSD of 1.04 Å^2^ (Fig. S8A). However, the active conformation of S1 that binds NAD^+^ contains an active-site cleft that is more closed relative to S1 in the B-oligomer (Fig. S8B). Specifically, α3 moves inwards towards the cleft, positioning the substrate-binding residues Tyr59 and Tyr63 for NAD^+^ binding (Fig. S8B) [39]. Since hu1B7 binds to α3, it could allosterically prevent the helix motion necessary to position Tyr59 and Tyr63 for substrate binding. Alternatively, hu1B7 could inhibit binding of the Gαi substrate. Since the loop containing S1 residues 112–127 is involved in Gαi binding in related toxins [39, 40], and is located on the opposite end of the active-site cleft as hu1B7 (Fig. S8C), this seems unlikely. However, if Gαi occupies a large volume in the active site cleft, hu1B7 could introduce steric clashes resulting in neutralization.

**Figure 5.**
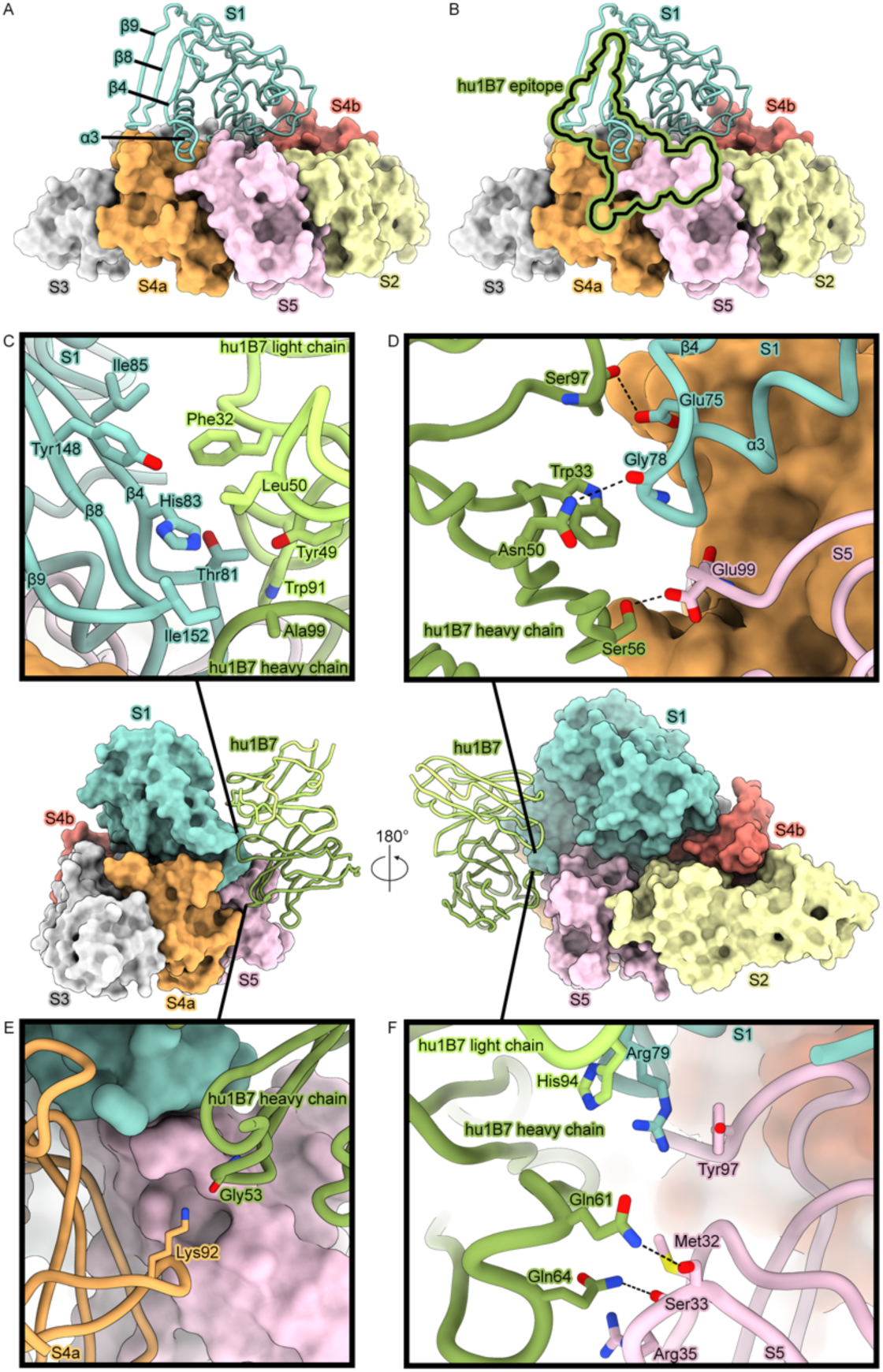
Hu1B7 binds a quaternary epitope spanning PTg S1 and S5. (A) PTg shown as molecular surfaces, with S1 colored turquoise, S2 colored yellow, S3 colored white, S4a colored orange, S4b colored red, S5 colored pink, hu11E6 (S2) colored blue, hu11E6 (S3) colored purple, and hu1B7 colored green. α3, β4, β8 and β9 secondary structural elements are labeled on S1. (B) Same as (A), with the outline of the hu1B7 epitope shown in black with a green outline. (C,D,E,F) Views of PTg bound to hu1B7 are shown in the middle row. Boxes show zoomed views depicting ribbon representations of hu1B7 and either S1 (C), S1 and S5 (D,F) or S4a (E), with key interface residues shown as sticks.

Unexpectedly, the hu1B7 heavy chain forms multiple contacts with PTg S5. Overall, the hu1B7 HC buries 262 Å^2^ and 166 Å^2^ on S1 and S5, respectively. hu1B7 HC Gly53 may also form a hydrogen bond with PTg S4a Lys92, but the map is not well-resolved in this region (Fig. 5E, Fig. S6), leaving the exact sidechain rotamer of PTg S4a Lys92 uncertain. Hu1B7 HC Gln61 and Gln64 form hydrogen bonds with the sidechain oxygen and mainchain oxygen, respectively, of S5 Ser33, with hu1B7 HC Gln64 packing onto S5 Met32 and S5 Arg35 (Fig. 5F). Hu1B7 LC His94 packs onto S1 Arg79, which itself forms a π–π stacking interaction with S5 Tyr97 as part of the interaction between S1 and the B-oligomer (Fig. 5F). While hu1B7 light chain His94 does not directly bind S5, it may stabilize the interaction between S1 Arg79 and S5 Tyr97. Binding of hu1B7 to both S1 and S5 suggests that hu1B7 binding may prevent S1 subunit dissociation from the B-oligomer. Although hu1B7 neutralization likely occurs via inhibition of PT retrograde trafficking [23], we previously hypothesized that clamping the S1 subunit to the B-oligomer via interactions with S5 could contribute to hu1B7 neutralization.

### Neutralization activity of hu1B7 mutants

To test whether clamping of S1 to S5 by hu1B7 contributes to the neutralization mechanism of hu1B7, we generated an hu1B7 variant (hu1B7_mut_) with point mutations expected to ablate the 3 hydrogen bonds between hu1B7 and S5: S56A, Q61A, Q64A. If S1–S5 clamping were critical to the hu1B7 neutralization mechanism, then ablation of these hydrogen bonds would prevent neutralization. However, hu1B7_mut_ binding to PT, as measured by ELISA, and PT neutralization of CHO cell clustering, were not detectably different from binding and neutralization by wild-type hu1B7 (Fig. S9). Therefore, S1–S5 clamping does not appear to play a role in hu1B7-mediated neutralization of PT.

## DISCUSSION

All acellular pertussis vaccines evaluated to date aim to induce high titers of neutralizing antibodies against PT since these mitigate the clinical symptoms of disease. This goal is complicated by the chemical detoxification of PT in most current vaccines, a process that appears to alter the structure of PT-neutralizing epitopes and their resulting immunogenicity. Characterization of key neutralizing epitopes on PT will contribute to a mechanistic understanding of PT protection and allow for their retention in future PT-based immunogens. Accordingly, we determined the structural basis for PT binding and neutralization by two synergistic and potently neutralizing antibodies: hu11E6 and hu1B7.

Antibody hu11E6 binds the B subunit to block PT-glycan interactions required for cell binding and toxicity. In the absence of antibodies, PT attachment to cells is mediated by the paralogous S2 and S3 subunits, which each contain two carbohydrate-binding domains: the N-terminal APT domain, which resembles mammalian C-type lectins, and the C-terminal AB_5_ toxin B-oligomer domain, which supports B subunit assembly [15, 41, 42]. Hu11E6 binds an APT domain epitope that is almost perfectly conserved across S2 and S3 (Fig. 3) and ablates PT interactions with sialylated N-glycans (Fig. 4B). This epitope surprisingly does not overlap with the structurally defined sialic acid-binding site in the C-terminal B- oligomer domain [13], nor does it affect binding to the sialyl Lewis^X^ blood group antigen or GQ2 ganglioside, which contain 1 and 4 terminal sialic acid residues, respectively (Fig 4D). Since the S2/S3 subunits engage four glycan classes [36], this suggests that the C-terminal B-oligomer domain is likely responsible for binding carbohydrates similar to Lewis^X^ and GQ2 and that other sites on PT may bind asialo-N-glycans. Overall, the N-terminal APT domain contains at least one binding site for a sialylated N-linked glycan moiety critical for PT cell adhesion whose interactions are blocked by hu11E6 binding.

Hu11E6 blocks PT attachment to mouse spleen and CHO cells [23, 43], prevents PT activation of human T cells (Fig. 4E) and is strongly protective in mouse infection models [31, 32, 44]. Mechanistic studies have shown that PT triggers immediate TCR-mediated intracellular signaling in T cells by binding an unknown receptor [37]. Here, we show that hu11E6 blocks this signaling and thus hu11E6-like antibodies may provide protection against PT-mediated T cell anergy and exhaustion. However, hu11E6 only blocks binding by a fraction of the glycan species engaged by PT (Fig 4A-D). Therefore, binding of sialylated N-glycans to the PT site blocked by hu11E6 represents a major attachment pathway *in vivo* but PT may engage different glycans to target other cell types as occurs in other AB_5_ toxins such as typhoid toxin [45]. The importance of hu11E6-like antibodies in preventing T cell activation bolsters the rationale for retaining B subunit epitopes in future PT immunogens.

Antibody hu1B7 prevents PT activity *in vitro* and potently protects infant baboons administered an at-birth dose from infection six weeks later [10]. The hu1B7 epitope on PT is dominated by S1 interactions, with ∼79% of the buried surface area on PT localized to S1 (Fig. 5). This is largely consistent with previous biochemical studies: hu1B7 contacts defining the hydrophobic interaction with S1 (L:W91, H:W33 and H: S97 with S1:H83 and Y148) and B-subunit interactions (L:H94 and S1:R79) were correctly identified (Fig S5D,E) [22, 38]. In addition, Sutherland *et al*., correctly predicted interactions with the B-oligomer based on decreased affinity of hu1B7 for recombinant S1 monomers.

However, the epitope was predicted to span the S1 and S4a subunits, whereas our structure shows the epitope spanning S1 and S5, with just one S4a contact (Fig. S5E). Antibodies binding ricin toxin epitopes that span the A-B toxin interface are also potently neutralizing [46], suggesting that this interface may be a general site of vulnerability.

Protection by hu1B7 is more likely related to altered intracellular trafficking than direct inhibition of catalytic activity. Access to the catalytic site is not impaired by hu1B7 binding although hu1B7- induced conformational changes could allosterically inhibit binding of NAD+ or the Gαi substrate (Fig. S8). Conversely, *in vitro* experiments with CHO cells suggested that hu1B7 alters PT intracellular trafficking to mediate neutralization [23]. As S1 must dissociate from the B-oligomer to reach its eventual target, one possibility is that hu1B7’s quaternary epitope clamps S1 to S5, frustrating S1 dissociation.

Accordingly, we simultaneously ablated the three hydrogen bonds formed between hu1B7 and PT S5. This had little effect on hu1B7 binding or neutralization (Fig. S9), suggesting that S1/S5 clamping is not relevant for the hu1B7 neutralization mechanism. A second possibility is that hu1B7 disrupts interactions required for PT retrograde trafficking. No mechanism has yet been described to mediate PT retrograde transport, but multiple mechanisms have been described for other AB toxins, including a C-terminal KDEL sequence for ER retention (cholera toxin), hitchhiking on a host protein that exhibits retrograde transport (shiga toxin) [47] and interactions with ER-resident chaperones [48]. Mutagenesis of the hu1B7- binding site could be used to generate non-binding probes to identify putative PT endosomal interaction partners to define this mechanism.

Despite the success of PT as a vaccine antigen, the chemical detoxification process used to inactivate PT in most approved aP formulations may have deleterious effects on antigenicity and carbohydrate binding. The engineered PTg used in this work [33] was rendered inactive by introducing R9K and E129G substitutions in the toxic S1 enzymatic subunit. Studies comparing PTd and PTg immunogenicity in humans observed higher total neutralizing titers and more durable responses after immunization with PTg-containing vaccines [28, 49, 50]; while sera from patients with confirmed pertussis infection contained higher titers of antibodies binding specific neutralizing epitopes (including the hu1B7 and hu11E6 epitopes) than sera from patients who recently received an acellular vaccination containing PTd [26, 27]. Interestingly, glutaraldehyde inactivation entirely ablated binding to a model sialylated N-glycan but only partially ablated binding to the asialylated version of the same carbohydrate [51]. The similar impacts of hu11E6 binding and glutaraldehyde inactivation on glycan binding suggest the same sialylated N-glycan site is affected. Therefore, intact carbohydrate sites on PTg likely represent important neutralizing epitopes that risk loss during chemical inactivation. Since the PT and PTg crystal structures are nearly identical (1.5 Å r.m.s.d.) [52], including in the hu1B7 and hu11E6 epitopes, this indicates PTg may elicit more effective immune responses by preserving key surface epitopes.

Defining the structural basis for potent PT neutralization by hu11E6 and hu1B7 helps to explain protection conferred by PT and will guide design of future PT immunogens. In addition, PT’s complex multi-subunit structure with unequal stoichiometry presents challenges for design of nucleic acid-based vaccines. Development of simpler PT-like immunogens may require retention of S5 sequences to elicit hu1B7-like antibodies and may benefit from focusing immune responses on the poorly antigenic S2/S3 subunits to elicit hu11E6-like antibodies. Together, this work supports the use of PTg over PTd in aP vaccines and suggests immunogen design strategies to elicit high titers of potently neutralizing antibodies.

## ACKNOWLEDGMENTS

We would like to thank Dr. Sasha Dickinson at the Sauer Structural Biology Laboratory at UT Austin for assistance with cryo-EM data collection. We thank Saowanee Suwanbenjakun and Wasin Buasri at BioNet-Asia for technical assistance with ELISA binding study. We acknowledge the University of Texas College of Natural Sciences and award RR160023 of the Cancer Prevention and Research Institute of Texas for support of the EM facility at the University of Texas at Austin. Glycan arrays were performed by the Protein-Glycan Interaction Resource of the CFG and the National Center for Functional Glycomics (NCFG) at Beth Israel Deaconess Medical Center, Harvard Medical School (supporting grant R24GM137763). This work was funded in part by Welch Foundation grant number F-0003-19620604 (J.S.M.) and National Institutes of Health R01AI155453 (J.A.M., J.S.M.).

## AUTHOR CONTRIBUTIONS

Conceptualization, J.A.G., J.A.M., and J.S.M.; Investigation and visualization, J.A.G., A.W.N.., R.E.W.; Critical reagents, W.W.; Writing – Original Draft, J.A.M., J.A.G.; Writing – Reviewing & Editing, J.A.G, A.W.N., R.E.W., W.W., J.A.M., and J.S.M.; Supervision, J.A.M. and J.S.M.

## METHODS

### Protein expression and purification

PTg, a genetically detoxified pertussis toxin containing R9K and E129G was produced from an engineered *Bp* BNA064 WCB strain [33]. This strain was cultured in supplemented MSS medium at 35°C in shake flask then transferred to a fermenter. After 32-48 h of growth, the culture supernatants were collected and processed with column chromatography to obtain a purified PTg and qualified using validated quality control tests. Full-length antibody versions of hu11E6 and hu1B7 were produced with human IgG1/ kappa constant domains as previously described [31]. To produce hu1B7 IgG and hu1B7 (hu1B7 S56A, Q61A, Q64A) IgG, FreeStyle 293 cells were co-transfected with hu1B7 heavy chain and hu1B7 light chain, or hu1B7 heavy chain S56A, Q61A, Q64A and hu1B7 light chain, respectively, using polyethylenimine (Polysciences). After 6 days, supernatant was harvested and IgG was purified using Protein A resin (Thermo Fisher). Fabs were generated by addition of 1:2000 wt/wt Endoproteinase Lys-C (Sigma) to hu11E6 IgG or hu1B7 IgG diluted to 1 mg/ml in PBS and incubated overnight at 37 °C. Subsequently, 1X Complete Protease Inhibitor Cocktail (Roche) was added to each reaction. The hu1B7 and hu11E6 Fabs were purified using CaptureSelect IgG-CH1 affinity resin (ThermoFisher) and further purified via size-exclusion chromatography using a Superdex 200 10/300 GL (Cytiva).

### Negative-stain electron microscopy

PTg at 0.04 mg/ml was mixed with a 2-fold molar excess of hu1B7 Fab and a 4-fold molar excess of hu11E6 Fab and allowed to bind for 20 minutes at room temperature in a glass tube. The complex (4.8 µl) was added to CU-CF400 grids (Electron Microscopy Sciences) that had been glow-discharged for 30s with using an Emitech K100X. The sample was then stained with NANO-W methylamine tungstate stain solution (NanoProbes). Micrographs were collected (50) using a 200 kV JEOL 2010F at a magnification of 120,000x (corresponding to 3.6 Å/pixel), at defocus values ranging from -0.5 μm to -1.5 μm. CTF correction, particle picking, and 2D class-averaging were performed in CryoSPARC.

### Cryo-electron microscopy

PTg (0.2 mg/ml in 20 mM Tris pH 7.5, 50 mM NaCl) was mixed with a 2- fold molar excess of hu1B7 Fab and a 4-fold molar excess of hu11E6 Fab and allowed to bind for 20 minutes at room temperature in a glass tube. The sample was then mixed in a 1:1 ratio with 20 mM Tris pH 7.5, 50 mM NaCl, 0.2% beta-octylglucoside resulting in a final concentration of 0.1 mg/ml PTg in 20 mM Tris pH 7.5, 50 mM NaCl, 0.1% β-octylglucoside. This sample (3 µl) was applied to an UltrAuFoil holey gold grid (Electron Microscopy Sciences) that was previously plasma-cleaned for 4 minutes using a Gatan Solarus 950 and then plunge-frozen using a Vitrobot Mark IV using +1 force, 3 second blot time, 100% humidity at 4 °C. Movies (1,500) were collected at a magnification of 22,500x with a Titan Krios and K3 detector, at a stage tilt of 30°. Motion correction, CTF estimation, particle picking, *ab-initio* reconstruction, heterogeneous refinement, and homogeneous refinement were performed in CryoSPARC. Map sharpening was performed using deepEMhancer. Model building and refinement were performed in Coot, Phenix, and ISOLDE. Cryo-EM data collection and refinement statistics are in Table S1. For the detergent screen, test cryo-EM images were taken in a 200 kV FEI Talos using an Ceta-M detector (ThermoFisher) (no detergent, 0.1% amphipol) or a 300 kV Titan Krios using an Ceta-M detector (ThermoFisher) (0.4% CHAPS, 0.04% fluoro-octyl maltoside, 0.1% β-octyl-glucoside).

### High-throughput glycan array

PTg at either 10 µM or 50 µM was incubated with 2-fold excess hu1B7 IgG (for detection) and either buffer or a 4-fold excess 11E6 Fab. The samples were then analyzed at the National Center for Functional Glycomics (NCFG) using version 5.5 of the NCFG glycan array containing 562 glycans. Each sample was analyzed in replicates of 6. The mean relative fluorescence unit (RFU) values for each array were analyzed in the NCFG’s GLAD software before being sorted into 3 categories: ‘sialylated N-glycan’, asialylated N-glycan’ and ‘other carbohydrate’ and plotted in GraphPad Prism 7.0.

### Binding assays

A high-binding 96-well ELISA plate (Costar) was incubated overnight at 4°C with 0.2 µg/mL PT (List Labs) in PBS, pH 7.4 or nothing (no coat). The plate was blocked with PBS-T-milk (PBS, pH 7.4 with 0.05% Tween-20 and 5% non-fat dry milk) for 1 hr at 25°C. Antibodies diluted to 5 µg/mL in PBS-T-milk were added to both the PT-coated and uncoated wells of the plate in duplicate and serially diluted 5-fold seven times. The antibodies were allowed to bind for 1 hr at 25°C. After washing with PBS-T (PBS, pH 7.4, with 0.05% Tween-20), a 1:2000 dilution of goat-anti-human Fc-HRP (Southern Biotech) in PBS-T-milk was added to each well and allowed to bind for 1 hr at 25°C. After washing, the plate was developed with TMB substrate (Thermo Fisher Scientific) and quenched with 1 N HCl. Absorbance was measured at 450 nm by using an Agilent BioTek Synergy H1 plate reader. Measurements were fit to a 4-parameter logistic curve.

### CHO cell clustering assay

Enzymatically active PT (List Labs) was diluted to 10 pM in CHO-K1 culture medium (high glucose DMEM with 10% fetal bovine serum and 1× penicillin and streptomycin). This PT diluent was used to prepare dilutions of antibodies hu1B7, hu1B7 S57A/Q62A/Q65A, or human IgG1 isotype control. The human IgG1 isotype control was also prepared in CHO-K1 culture medium without pertussis toxin (Isotype Control – no PT). The highest concentration of each antibody was 200 nM, with seven additional 2-fold dilutions prepared in wells of a black-walled, clear bottom 96-well plate (Agilent). The PT and antibody mixtures were incubated at 37°C for 30 min, then an equal volume of CHO-K1 culture medium with 10^4^ CHO-K1 cells per well were added to the plate, diluting the pertussis toxin and antibodies by 2-fold. The plate was allowed to incubate at 37°C for approximately 40 hr, then washed with PBS+MgCa (PBS containing magnesium and calcium chloride, Sigma). PBS+MgCa with 1 µg/mL Hoescht 33342 nuclear stain and 1 µM CFSE cytoplasmic stain was added to each well and the plate was incubated at 37°C for 15 min. The plate was then washed one time with PBS+MgCa, one time with CHO- K1 culture medium, two times with PBS+MgCa, then 50 uL of PBS+MgCa was added to each well for imaging. The plates were imaged with a Cytation C10 plate imager using a 10x objective capturing fluorescent images at 447 nm with 377 nm excitation (Hoescht 33342) and 525 nm with 469 nm excitation (CFSE). Four images were captured and a montage created for each well. Masking was applied to measure the area of non-CFSE fluorescent regions greater than 100 nm in diameter in each well. The total empty area of each well was transformed to the Normalized Clustering Score by subtracting the average empty area of the 100 nM Isotype Control – no PT from each empty area value and dividing the result by the subtracted average empty area of the 100 nM Isotype Control with PT. These final data were fit to [Inhibitor] vs. response equations using Prism (GraphPad).

### T cell activation assay

PBMCs from a healthy anonymized donor were isolated from 50 mL Leukopak (Gulf Coast Regional Blood Center). Briefly, blood was diluted with PBS (1:1) and overlaid over Corning Lymphocyte Separation Medium (Cat 25072CV). This was centrifuged for 30 min at 400g (no brake), the lymphocyte layer was carefully removed, and residual red blood cells were lysed with RBC lysis buffer (ThermoFisher Cat 00-4333-57). Cells were washed with PBS and cultured in 500 mL RPMI (Sigma Cat R8758) with 50 mL heat inactivated FBS (Gibco Cat A52567-01), penicillin-streptomycin (Sigma Cat P4458), 5 mL 100X non-essential amino acids (Gibco Cat 11140-050), 5 mL 100X GlutaMAX (Gibco Cat 35050-061), 5 mL of 100 mM sodium pyruvate (Gibco 11360-070; PBMC media). PTg (1 μg/ml = 9.5 nM final concentration) was incubated with serially diluted Fab in 100 μl PBMC media for 1 hr at 37 °C, then combined with an equal volume of PBMCs (5 x 10^5^ per well) and co-incubated overnight. Cells were washed twice with 1% FBS in PBS (staining buffer) before incubation with 2.5 μg Fc block (BD Biosciences Cat 564219) in staining buffer for 15 min at room temperature. Staining buffer containing 1:250 concentration of anti-CD3-PE, anti-CD69-APC, anti-CD4-FITC, and anti-CD8-BV421 (Biolegend Cat 317308, 310910, 317408, 344747) was added directly to an equal volume of the Fc block and incubated at room temperature for 20 minutes. Cells were washed twice with staining buffer before collecting 10,000 events were collected per sample on a ThermoFisher Attune Flow Cytometer.

Sequential gating was performed based on size (FSC vs SSC), single cells (FSC-A vs FSC-H), the CD3- expressing population and finally CD69-expression based on untreated negative controls. Data were plotted using FlowJo 10.7.1 and Graphpad Prism 9.50; these data were fit to [inhibitor] vs. response equations to determine IC_50_ values. Samples were run in duplicate with two experimental repeats per donor.

