## Supplemental figures for "Structural Basis for Antibody Neutralization of Pertussis Toxin"

**Affiliations:**

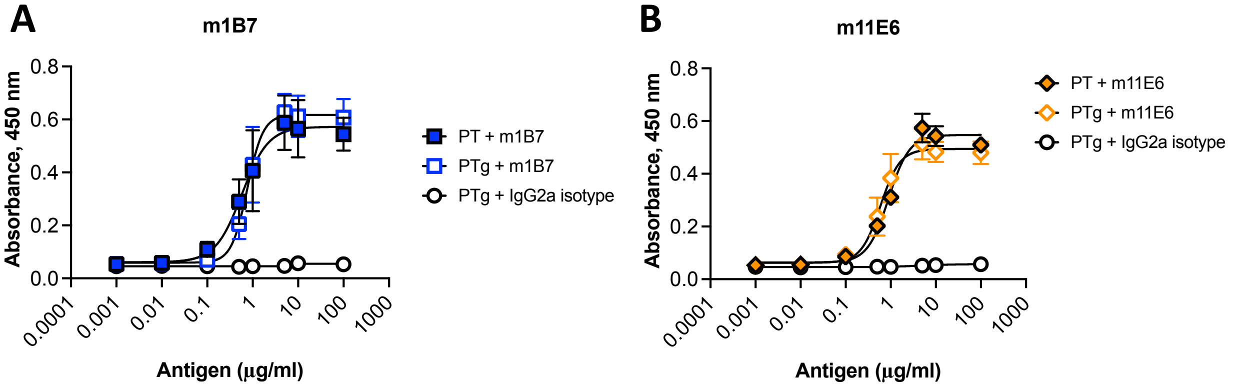


**Figure S1. Antibodies bind PT and PTg similarly.** Indirect ELISA was used to compare binding to coated PT and PTg.

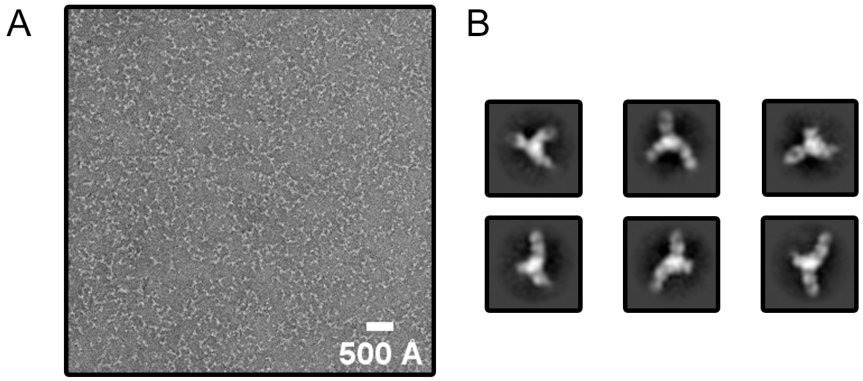


**Figure S2. Negative-stain electron microscopy of PTg in complex with hu11E6 and hu1B7**

(A) Negative-stain electron microscopy micrograph of PTg in complex with hu11E6 and hu1B7. (B) Negative-stain electron microscopy 2D class-averages of PTg in complex with hu11E6 and hu1B7.

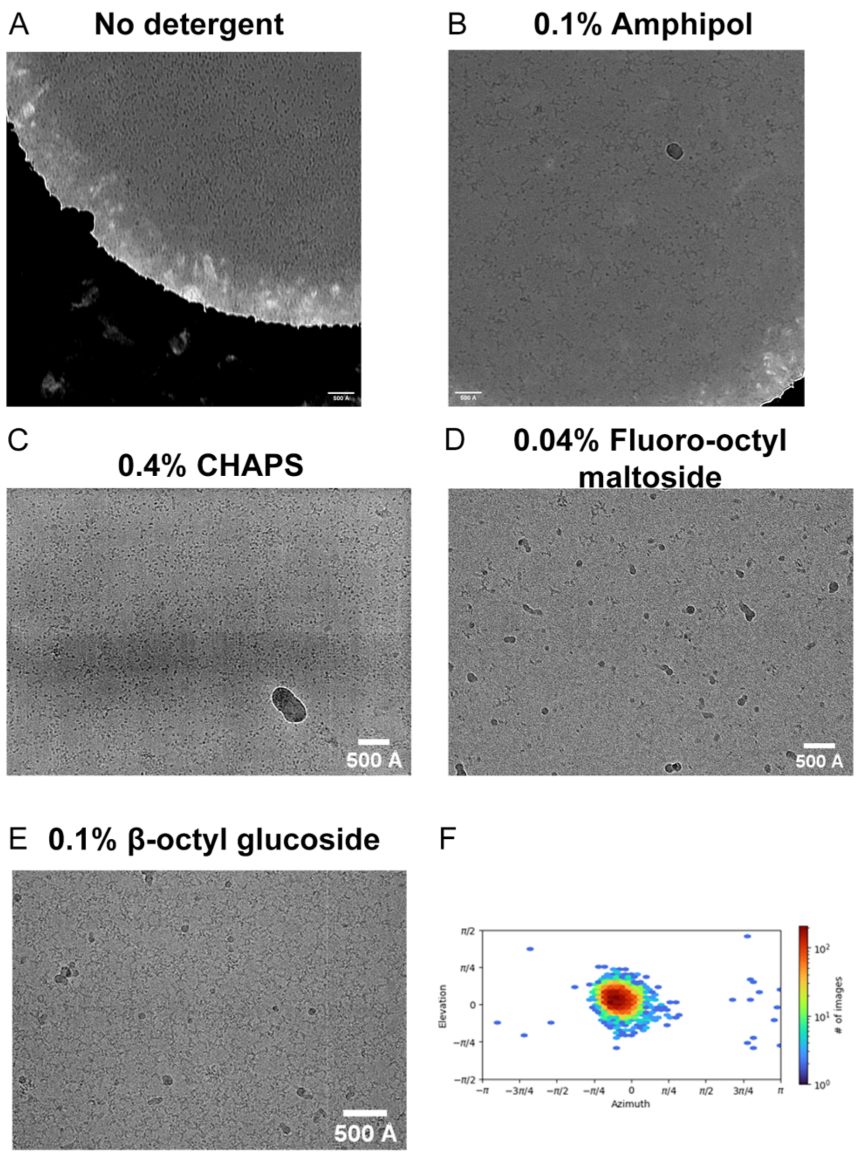


**Figure S3. Screening of cryo-electron microscopy freezing and collection conditions**

(A-E) Representative screening micrographs of cryo-electron microscopy grids containing (A) no detergent, (B) 0.1 % amphipol, (C) 0.4% CHAPS, (D) 0.04% fluoro-octyl-maltoside, and (E) 0.1% β-octylglucoside. (F) Orientation distribution plot from a homogeneous refinement of particles extracted from a test dataset taken of the grid in (E) containing 0.1% β-octylglucoside.

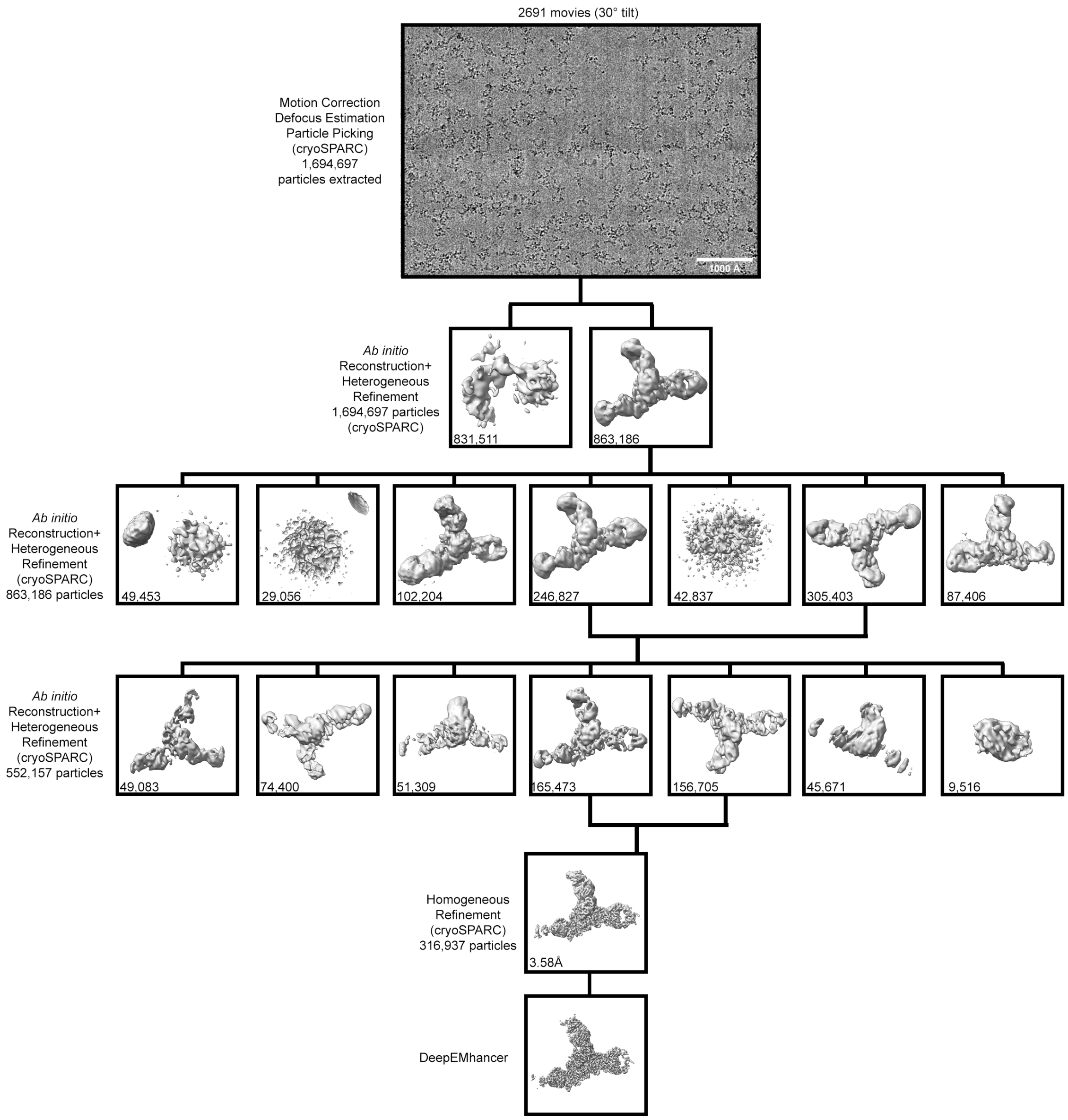


**Figure S4. Cryo-electron microscopy data processing workflow**

Cryo-electron microscopy data processing workflow showing the volumes resulting from 3 rounds of heterogeneous refinement of extracted particles followed by homogeneous refinement of the final particle stack. The number of final particles in each heterogeneous refinement class is shown in the box with its corresponding volume.

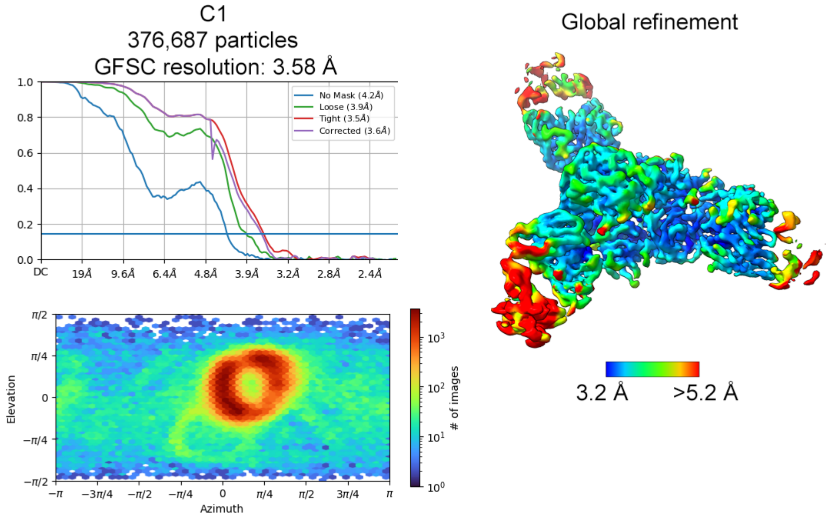


**Figure S5. Cryo-electron microscopy data processing validation**

Left: Fourier shell correlation curve and view distribution plot for the cryo-EM map of PTg in complex with hu11E6 and hu1B7. Right: cryo-EM map of PTg in complex with hu11E6 and hu1B7 colored by estimated local resolution.

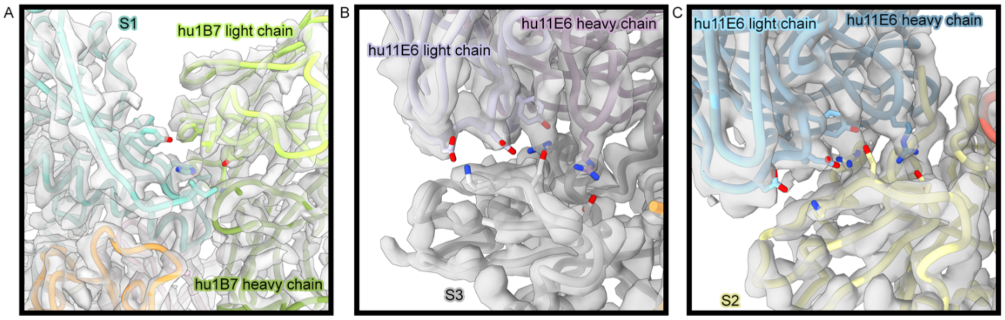


**Figure S6. Validation of model in cryo-EM map**

Example images of the model of PTg in complex with hu11E6 and hu1B7 with the cryo-EM map from the interface of 1B7 and PTg (A), 11E6 and S3 (B) and 11E6 and S2 (C).

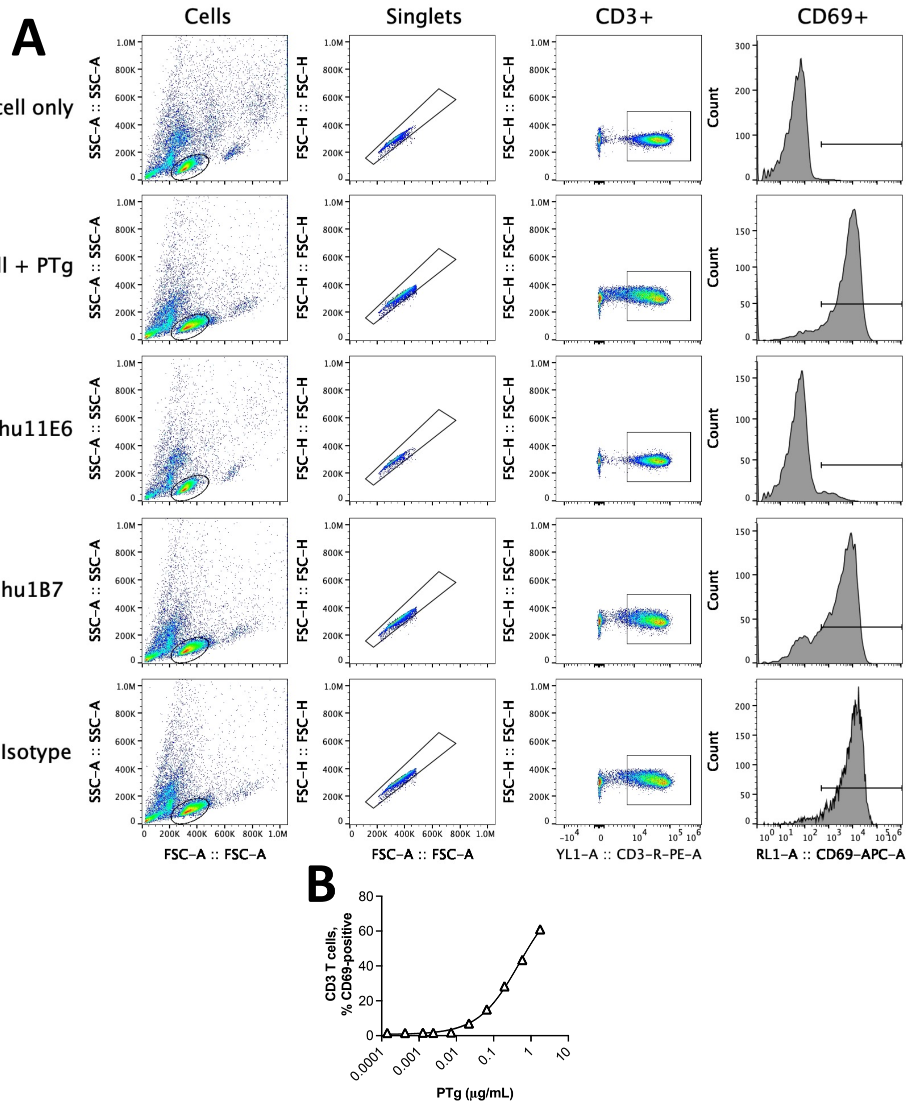


**Figure S7. Gating strategy for PTg activation of primary T cells**

(A) Example gating strategy to isolate activated T cells. PBMCs are first gated by size (FSC v SSC), singlet discrimination (FSC-A v FSC-H), and CD3+ cells before scanning for CD69 upregulation. Data shown are from the sample with the highest concentration of antibody for each condition including an antibody. (B) Human donor PBMCs were coincubated with various amounts of PTg for 24 hours before analysis for CD69 upregulation of CD3+ T cells. Data shown are representative of two biological replicates, with each point showing the average of two technical replicates plus/ minus the observed range.

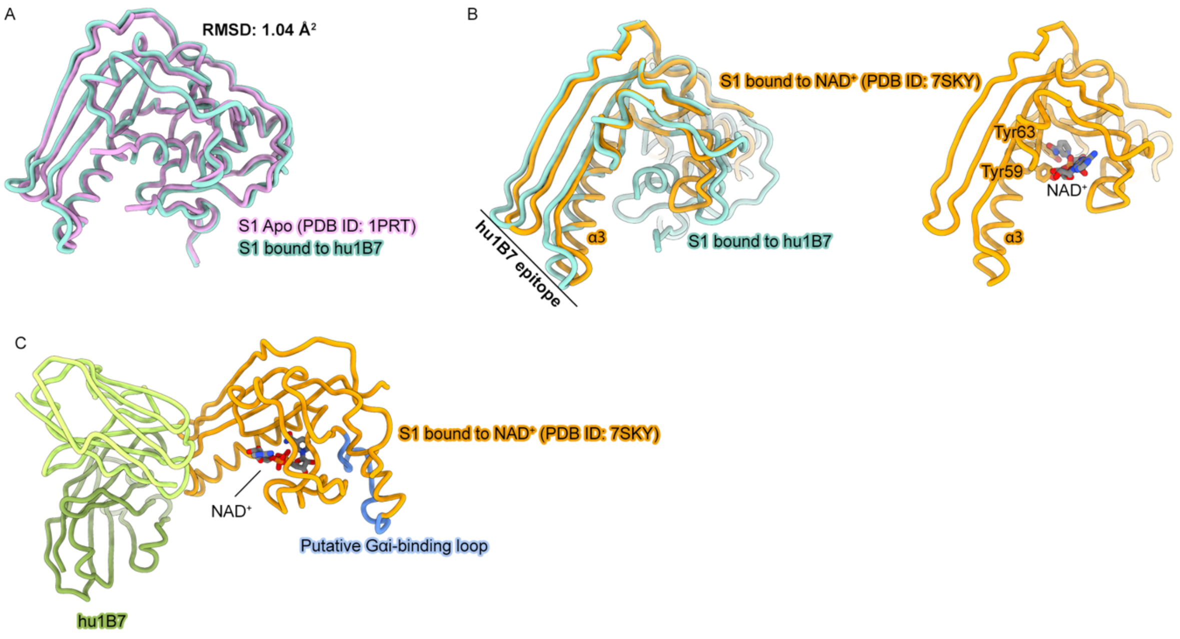


**Figure S8. Structural analysis of catalytic and substrate-binding sites in PTg bound to 1B7**

(a) PTg S1 bound to 1B7 (cyan) superimposed with apo PT S1 (violet, PDB ID: 1PRT). (B) Left: PTg S1 bound to 1B7 (cyan) superimposed with NAD+-bound PT S1 (orange, PDB ID: 7SKY). Right: NAD+-bound PT S1 (orange) with NAD+ shown as sticks (PDB ID: 7SKY). (C) 1B7 Fab (green) modeled bound to to NAD+-bound PT S1 (orange, PDB ID: 7SKY) with NAD+ shown as sticks and the putative $G\alpha_i$-binding loop of S1 colored blue.

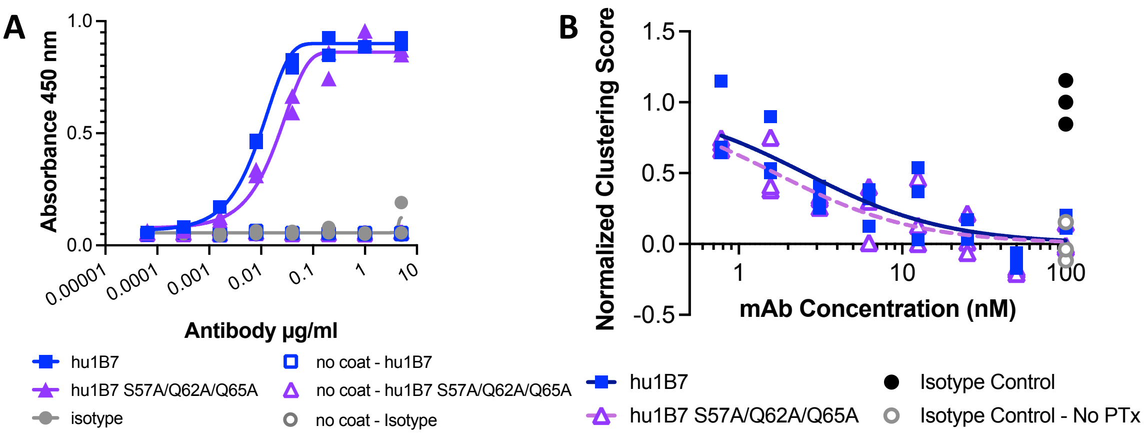


**Figure S9. Binding and neutralization activity is retained in hu1B7 variants with ablated S5 binding**

(A) ELISA binding of hu1B7 (blue squares), hu1B7 variant S57A/Q62A/Q65A which should disrupt binding to the S5 subunit of PTx (magenta triangles), or an isotype control antibody (grey circles) to either PT coated (solid lines, filled symbols) or no coat (dotted lines, open symbols) wells. (B) CHO cell clustering measurements of CHO-K1 cells incubated with no PT (grey circles), 5 pM PT and 100 nM isotype control antibody (black diamonds), 5 pM PT and hu1B7 (blue squares) or hu1B7 variant S57A/Q62A/Q65A (magenta triangles) concentrations ranging from 780 pM to 100 nM in 2-fold increments. Data shown are representative of two replicate experiments.

### Table S1. Cryo-EM data collection and refinement statistics

| **EM data collection** | **Pertussis Toxin+hu1B7 Fab+hu11E6 Fab** |
|---|---|
| Microscope | Krios |
| Detector | K3 |
| Defocus range (μm) | 0.8–2.2 |
| Tilt angle (°) | 30 |
| Pixel size (Å) | 1.1 |
| Magnification | 22,500x |
| Voltage (kV) | 300 |
| Exposure rate ($e^{-}$/pix/sec) | 8 |
| Frames per exposure | 80 |
| Micrographs collected | 1,503 |
| Micrographs used | 1,500 |
| Electron exposure ($e^{-}$/Å$^{2}$) | 49 |
| Particles extracted | 1,694,697 |
| Automation software | SerialEM |
| **3D reconstruction statistics** | |
| Final particles | 316,937 |
| Symmetry | C1 |
| Unmasked resolution at 0.5 FSC (Å) | 7.9 |
| Masked resolution at 0.5 FSC (Å) | 4.1 |
| Unmasked resolution at 0.143 FSC (Å) | 4.2 |
| Masked resolution at 0.143 FSC (Å) | 3.5 |
| **Model refinement and validation** | |
| Refinement package | Phenix |
| Refinement tool | Real-space refinement |
| Refinement strategies | min global, local_grid_search, adp, ss restraints, rotamer restraints, Ramachandran restraints |
| Initial model(s) used (PDB ID) | 1PRT |
| Model composition | |
| Number of atoms | 12455 |
| Protein B-factors (mean) | 60.5 |
| RMSD bond lengths (Å) | 0.002 |
| RMSD bond angles (°) | 0.56 |
| MolProbity score | 1.49 |
| Clashscore | 4.6 |
| Rotamer outliers (%) | 0.07 |
| C-beta outliers (%) | 0.0 |
| Ramachandran plot | |
| Favored (%) | 96.2 |
| Allowed (%) | 3.8 |
| Outliers (%) | 0.0 |
| EMRinger score | 3.27 |
| CaBLAM outliers (%) | 2.3 |
| CC (mask) | 0.74 |
| **Data Availability** | |
| EMDB | EMD-XXXXX |
| PDB | XXXX |
